# Influ-BERT: A Domain-Adaptive Genomic Language Model for Advancing Influenza A Virus Research

**DOI:** 10.1101/2025.07.31.667841

**Authors:** Rongye Ye, Lun Li, Ana Tereza Ribeiro Vasconcelos, Shuhui Song

## Abstract

Influenza A Virus (IAV) poses a persistent threat to global public health due to its broad host adaptability, frequent antigenic variation, and potential for cross-species transmission. Accurate identification of IAV subtypes is essential for effective epidemic surveillance and precise disease control. Here, we present Influ-BERT, a domain-adaptive pretrained model based on the Transformer architecture. Optimized from DNABERT-2, Influ-BERT was developed using a dedicated corpus of approximately 900,000 influenza genome sequences. We constructed a custom Byte Pair Encoding (BPE) tokenizer, and employed a two-stage training strategy involving domain-adaptive pretraining followed by task-specific fine-tuning. This approach significantly enhanced identification performance for IAV subtypes. Experimental results demonstrate that Influ-BERT outperforms both traditional machine learning approaches and general genomic language models, such as DNABERT-2 and MegaDNA, in the task of IAV subtype identification. The model achieved F1-scores consistently above 97% and exhibited stable performance gains for subtypes that are underrepresented in sequencing data, including H5N8, H5N1, H7N9, and H9N2. Beyond subtype identification, Influ-BERT was successfully applied to additional tasks including respiratory virus identification, IAV pathogenicity prediction, and identification of IAV genomic fragments and functional genes, demonstrating robust performance throughout. Further interpretability analysis using sliding window perturbation confirmed that the model focuses on biologically significant genomic regions, providing insight into its improved predictive capability.

**Contact:** (Song S), (Ana Tereza Ribeiro Vasconcelos)

## Introduction

Influenza viruses are among the most epidemiologically significant respiratory pathogens globally, posing a long-standing and significant threat to global public health [1,2]. Among these, Influenza A viruses (IAV) constitute the primary pathogen responsible for human influenza pandemics due to its broad host range and capacity for antigenic variation. IAV is an RNA virus, and its genome is composed of eight single-stranded negative-sense RNA segments [3]. Specifically, Segment 4 encodes hemagglutinin (HA), and Segment 6 encodes neuraminidase (NA). These two surface glycoproteins determine viral subtype characteristics; they mediate viral entry into and release from host cells and are the primary targets for the host immune system. The generation of new HA-NA combinations through gene reassortment provides the molecular basis for cross-species transmission and pandemics by IAV [4]. For instance, the highly pathogenic avian influenza virus of the H5N1 subtype is one of the most significant pandemic threats currently identified. They have demonstrated the capacity for transmission among multiple mammalian species, indicating a potential for cross-species transmission to humans [5]. The widespread prevalence of H9N2 in live bird markets increases the zoonotic risk [6]. Similarly, H7N9, an avian influenza virus that can be highly pathogenic in poultry, poses a considerable threat to the poultry industry and public health, with human infections associated with a high mortality rate [7].

Traditional IAV subtype detection mainly relies on molecular biology techniques like reverse transcription PCR (RT-PCR) for accurate identification of common subtypes [8,9]. Serological methods, including hemagglutination inhibition (HAI) assays and neutralization assays, are also widely employed. HAI detects subtype-specific antibodies by their ability to inhibit hemagglutination [10], while neutralization assays identify subtypes by measuring antibody-mediated neutralization of viral infectivity [11]. However, these conventional methods have limited throughput, posing challenges for large-scale automated sample analysis. Advances in high-throughput sequencing have enabled genome-based methods for influenza subtyping, primarily through sequence alignment against reference databases [12]. These approaches recognize subtypes based on sequence similarity and facilitate the analysis of evolutionary relationships through phylogenetic trees [13,14]. Tools like NextClade [15] are particularly valuable for detecting novel subtypes or recombinant variants. However, alignment-based strategies face two key challenges: the high mutation rate IAV requires constant database updates, rendering pre-built references rapidly obsolete, and the computational complexity of alignment and phylogenetic tree construction becomes prohibitive for large-scale analyses.

The advent of artificial intelligence has transformed influenza subtype identification, with machine learning methods increasingly applied for this purpose. Current approaches typically use k-mer frequencies as features to train identification models [16-18]. However, the performance of these methods is highly dependent on manually designed features, such as the selection of k-mer length, making them sensitive to the quality feature extraction. Furthermore, these conventional models are limited in their ability to capture contextual dependencies within viral genome sequences. Recent advances in Transformer architecture have created new opportunities for biological sequence analysis [19-22]. Leveraging the symbolic similarity between nucleic acid sequences and natural language, researchers have developed numerous pretrained genomic language models [23,24], such as DNABERT [25,26], MegaDNA [27], and RNA-specific models like LucaProt [28], which integrating RdRp amino acid sequences with predicted protein structures in a multimodal Transformer framework, thereby enhancing virus discovery and the exploration of viral “dark matter.”

However, long-tail data distribution is prevalent across various real-world applications. This distribution pattern is characterized by a severe class imbalance, where most samples are concentrated in a small number of categories, while a large number of remaining categories contain relatively few samples. In the context of IAV, subtypes that have demonstrated sustained and widespread transmission among humans (e.g., H1N1 and H3N2) are classified as frequent subtypes. In our dataset, subtypes with an occurrence frequency below 5% and only limited documented human infection cases were classified as low-frequency subtypes. Due to the long-tail distribution of IAV data, models exhibit strong identification capabilities for prevalent subtypes (H1N1, H3N2), but performance declined significantly for low-frequency subtypes. These low-frequency subtypes remain highly important for epidemiological surveillance [29-31]. Consequently, the development of robust subtype prediction models is highly valuable for early warning and prevention for influenza pandemics.

Here, we propose Influ-BERT, a model inspired by DNABERT-2 architecture, that was specifically optimized to enhance identification capabilities for low-frequency IAV subtypes. We evaluated Influ-BERT against several baseline approaches, including k-mer-based machine learning methods (Random Forest, SVM, Logistic Regression), the masked pretrained model DNABERT-2, and the generative model MegaDNA. The results demonstrate that Influ-BERT outperforms these existing methods in identification low-frequency subtypes while maintaining strong overall performance. Moreover, to extend the applicability of Influ-BERT, we additionally designed three biologically meaningful downstream tasks: identification of common respiratory viruses (including SARS-CoV-2, rhinovirus, respiratory syncytial virus, and influenza virus), identification of influenza virus genomic fragments and functional genes, and prediction of IAV pathogenicity. Experimental results indicate that Influ-BERT achieves superior performance across these tasks, demonstrating its broad utility in influenza genome analysis. Finally, we introduced sequence perturbation analysis based on a sliding window approach, which revealed differences in feature identification patterns between Influ-BERT and DNABERT-2, providing a preliminary explanation for the performance improvement of Influ-BERT’s.

## Materials and Methods

### Dataset

We systematically collected approximately 900,000 influenza virus genome sequences from the NCBI Nucleotide database [32], encompassing more than 100 naturally occurring HA–NA combinations that have been detected to date. The dataset comprises approximately 800,000 IAV sequences, ∼100,000 influenza B virus (IBV) sequences, and over 4,000 influenza C and D virus (ICV and IDV) sequences. This dataset was utilized in two key steps: (1) the construction of a BPE vocabulary specific to influenza virus sequences; and (2) domain-adaptive pretraining of the language model. To evaluate the performance of the adapted model comprehensively, we designed four downstream tasks with biological relevance.

Task1 focuses on subtype identification. We first established an identification workflow that is agnostic to specific genomic fragments, encompassing tasks for five and ten subtypes. This workflow is particularly suited for rapid detection scenarios and cases with limited sequencing coverage. Considering the requirements of high-precision applications, such as vaccine design, which necessitate the analysis of specific influenza genomic segments, we further developed a dedicated training and subtyping pipeline focused exclusively on hemagglutinin (HA) and neuraminidase (NA) fragments. For this task, we utilized a dataset of 40,000 independently collected IAV sequences representing 66 subtype combinations, which were excluded from the domain-adaptive pretraining corpus. The dataset exhibits a long-tail distribution (**Figure 1a**), and a length distribution of sequences within the domain-adaptive pretraining corpus is presented in **Figure 1b**. For the subtype identification workflow relying exclusively on HA and NA fragments, we extracted the corresponding HA and NA segment data from these 40,000 sequences. The distribution of different genomic fragments is shown in **Figure 1c**.

**Figure 1.**
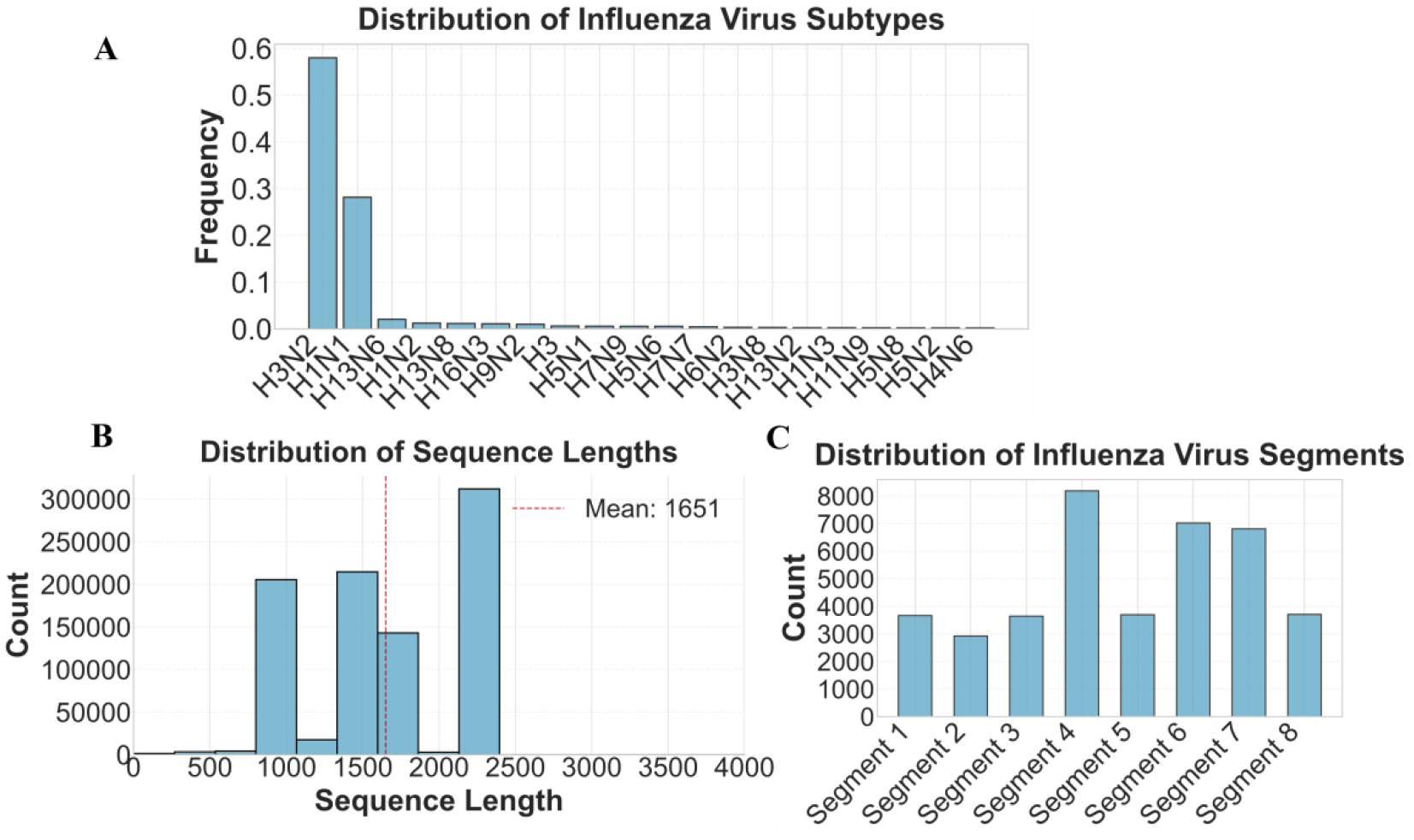
Statistical distribution of data. a) The frequency distribution of the top 20 subtypes in the down-stream task dataset; b) Length distribution of segments within the corpus used during the domain-adaptive pretraining phase; c) Distribution of the number of different genomic segments across downstream task datasets.

Task2 is designed to identify common respiratory viruses, including SARS-CoV-2, rhinovirus, respiratory syncytial virus (RSV), and influenza virus. Approximately 10,000 sequences were selected for each virus. For every virus type, the corresponding genomic fragments were extracted. To mitigate potential biases arising from variations in sequence length, all viral fragments were constrained to a maximum length of 10,000 base pairs.

Task3 focuses on the identification of IAV genomic fragments and functional genes. For each accession in the Task1 dataset, the corresponding fragment labels, functional gene sequences, and their associated annotations were retrieved from the NCBI database.

Task4 is dedicated to predicting the pathogenicity of IAV sequences. We constructed a label mapping dictionary to define pathogenicity categories (see Supplementary Table S1) and assembled a dataset of 1,662 sequences with relatively well-defined pathogenicity categories. To address the common challenge of limited data availability in real-world scenarios, this task specifically evaluates model performance under data-scarce conditions. Accordingly, we designed two limited-data learning experiments to systematically evaluate predictive performance: one trained on 200 samples (100 high- and 100 low-pathogenicity) and the other on 400 samples (200 high- and 200 low-pathogenicity), with the remaining sequences reserved for independent testing.

With the exception of Task4, all datasets were partitioned into training, validation, and test sets using a ratio of 8:0.5:1.5. Model performance was evaluated using accuracy, precision, recall, and F1-score as the primary metrics, providing a robust quantitative assessment.

### Experiment Setting

#### Software Configuration

During the domain-adaptive pretraining phase, BPE vocabulary with a size of 512 was constructed. All other training hyperparameters were kept consistent with those used in DNABERT-2. All experiments were conducted using the PyTorch framework (version 2.5.1), with model training and evaluation implemented via the Hugging Face Transformers library (version 4.49.0).

#### Baseline Model

This experiment compares five models, including three machine learning models based on k-mer (3-mer) encoding, the DNABERT-2 model based on masked pretraining, and the MegaDNA model based on generative pretraining: 1) **Random Forest (RF) [33]**: An ensemble learning model that improves model generalization by combining the predictions of multiple decision trees. It is widely used in classification and regression tasks; 2) **Support Vector Machine (SVM) [34]**: A supervised discriminative model whose core goal is to find an optimal hyperplane that maximizes the separation between samples of different classes; 3) **Logistic Regression (LR) [35]**: A commonly used classification machine learning model for classification that outputs a probability representing the likelihood of a sample belonging to a certain class. It is widely applied in classification tasks; 4) **DNABERT-2 [26]**: A self-supervised pretrained DNA language model that uses BPE encoding. It was pretrained on a large-scale multi-genome dataset containing 32.5 billion nucleotide bases and covering 135 species. This model can capture complex dependencies in DNA sequences and has demonstrated superior performance in multiple biological tasks; 5) **MegaDNA [27]**: A generative pretrained long-sequence DNA model that uses nucleotide-level tokenization. This model can handle genomic sequences up to 96,000 base pairs and has the capability for sequence generation and classification of unannotated sequences.

### Mask Language Model

Inspired by the success of natural language processing (NLP), masked pretraining has emerged as an effective paradigm for learning representations from genomic sequences. As a classical self-supervised learning strategy, masked pretraining operates by randomly masking a subset of input tokens during training and requiring the model to predict the original tokens based on their surrounding context. This approach enables the model to capture deep semantic representations of sequences. The objective function is designed to maximize the likelihood of the masked tokens, which is formally defined as follows:

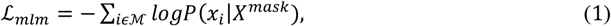

where *x*_*i*_ represents the token at position *i* in the original sequence, and *X*^*mask*^ refers to the masked input sequence. *P*(*x*_*i*_|*X*^*mask*^) denotes the model’s probability of reconstructing *x*_*i*_ given the masked input *X*^*mask*^. The Influ-BERT model adopts the same architectural framework as DNABERT-2, utilizing the Transformer attention mechanism as its core component for sequence representation learning. The attention function [14] is computed as follows:

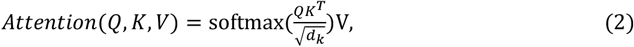

where *Q* denotes the query matrix, representing the current input query; *K* denotes the key matrix, which typically encodes the features of candidate elements; *V* is the value matrix, containing the information associated with each key; and *d*_*k*_ is a scaling factor used to normalize the dot product of queries and keys. Within the Transformer architecture, knowledge representations across different subspaces are typically learned using a multi-head attention mechanism. The computation is defined as follows:

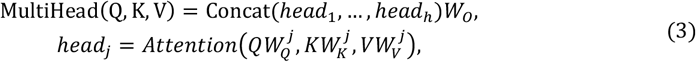

where *W* denotes the learnable projection matrices applied during linear transformations, and *head*_*j*_ refers to the *j*-th attention head among a total of *h* heads in the multi-head attention mechanism. Finally, during the fine-tuning phase, the task-specific loss function is defined as the cross-entropy loss, which is formulated as follows:

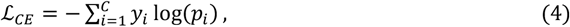

where *C* denotes the total number of classes, *y*_*i*_ represents the ground-truth label for the sample, and *p*_*i*_ denotes the predicted probability that the sample belongs to class *i*.

### Transfer Knowledge from Domain-Adaptive Model

Although genome language models pretrained on cross-species genomic data have achieved remarkable performance across a variety of biological sequence analysis tasks, directly applying them to influenza virus genotyping remains a significant challenge. The intrinsic characteristics of influenza viruses—namely, their long-tailed data distribution and high mutation rates constrain the fine-tuning performance of models such as DNABERT-2. To address this limitation, we first adopt DNABERT-2 as the foundational architecture, which has been pretrained on a large-scale dataset comprising 32.5 billion bases from 135 species, including bacteria, fungi, and mammals. Subsequently, we train a BPE tokenizer using an influenza-specific corpus and perform domain-adaptive pretraining on this corpus to enhance the model’s capacity for representing influenza viral sequences. We set the vocabulary size from 4096 to 512, while keeping all other pretraining hyperparameters consistent with those of DNABERT-2. For model adaptation, we employ a two-stage learning strategy: (1) domain-adaptive pretraining to enable the model to learn general representations of influenza sequences; and (2) transfer learning by fine-tuning the domain-adaptive pretraining model on downstream tasks with the addition of task-specific classification heads. During fine-tuning, we apply a supervised learning strategy with full parameter updates on Influenza sequences, optimizing the model using the cross-entropy loss function (Equation 4). This approach effectively combines the model’s ability to capture general sequence features of influenza viruses with task-specific learning [36], thereby improving its performance across diverse downstream tasks.

### Sequence Perturbation Analysis

The core idea of sequence perturbation analysis is to systematically mask local regions of the input sequence and then feed the masked sequences into the model to observe the resulting changes in output. This allows for a quantitative assessment of each sequence fragment’s contribution to the model’s decision. Specifically, for an influenza viral genome sequence *S* = (*s*_1_, *s*_2_, …, *s*_*n*_), and a set of class labels *C* = {*c*_1_, *c*_2_, …, *c*_*m*_}, we define a window size *w*, and first compute the model’s prediction on the original input sequence *S*:

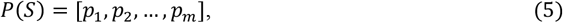

where 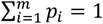. Subsequently, for a window starting at position *j*, we set the *w* = 5, and apply masking to this region. The perturbed sequence is defined as:

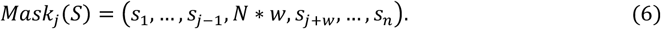

Here, *N* denotes the masking operation. Finally, for a sequence labeled as class *c*, the importance score of the window is computed as:

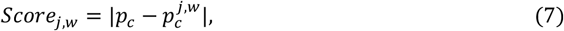

where *p*_*c*_ denotes the model-predicted probability of class *c* for the original input sequence, and 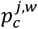 denotes the predicted probability of class *c* after masking a window of size *w* starting at position *j* in the original sequence.

## Results

### Overview of Influ-BERT

As shown in **Figure 2**, the Influ-BERT framework comprises by the domain-adaptive pretraining module and sequence recognition fine-tuning module. We first construct a vocabulary tailored for an influenza-specific corpus sequences using the BPE algorithm. Then, all influenza virus sequences in the corpus were tokenized using this vocabulary. The pre-trained model was initialized with the DNABERT-2 pretrained weights and conducted for a domain adaptive pretraining using masked language model (MLM) objective. For different downstream tasks, we constructed three datasets and consistently applied the influenza-specific BPE vocabulary to tokenize the input sequences. Following a transfer learning paradigm, we leveraged the domain-adaptive model to extract features, upon which task-specific classification heads were added to enable end-to-end fine-tuning. Specifically, we designed four biologically meaningful downstream tasks: (1) improving the subtyping accuracy for low-frequency IAV subtypes; (2) identifying common respiratory viruses; (3) recognizing IAV genomic segments and functional genes; and (4) predicting IAV pathogenicity. A sliding window–based perturbation analysis was further conducted to provide preliminary interpretability of the model’s predictions.

**Figure 2.**
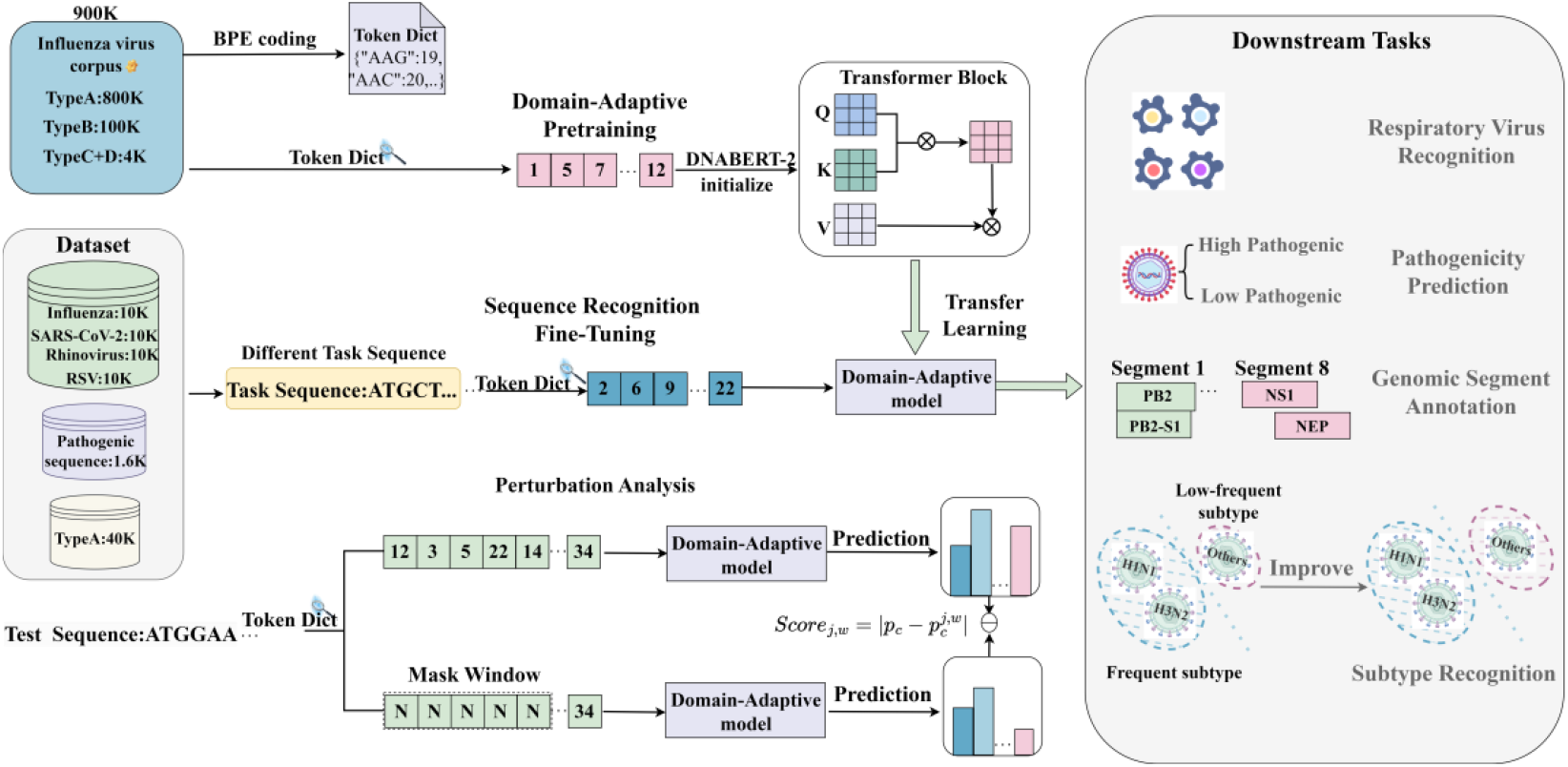
Schematic overview of the Influ-BERT framework. a) A BPE tokenizer is trained on a curated influenza-specific corpus to construct a domain-specific vocabulary, which is subsequently used for domain-adaptive pretraining, based on the DNABERT-2 architecture; b) Sequence recognition fine-tuning workflow: After domain-adaptive pretraining, downstream tasks are performed via transfer learning combined with task-specific fine-tuning; c) Interpretability analysis: A masking-based perturbation strategy is employed to identify attention-sensitive regions, enabling interpretation of model predictions.

### Performance Evaluation for IAV Subtype Identification

We systematically evaluated the performance of Influ-BERT on the task of IAV subtype identification without distinguishing genomic segments, encompassing high-frequency subtypes (H3N2 and H1N1) as well as low-frequency subtypes (frequency < 5%). This scenario is particularly relevant for rapid detection and situations with insufficient sequencing coverage. Among these low-frequency subtypes, H5N1, H7N9, and H9N2, with frequencies 0.6%, 0.5%, and 1%, were prioritized for analysis due to their relatively significant impact on public health as reported in previous studies. We assessed performance using four metrics—accuracy, precision, recall, and F1-score, and found that Influ-BERT outperformed all other models across overall dataset performance metrics (As shown in **Table 1**).

**Table 1.** Influ-BERT was compared with baseline models on subtype recognition.

| Task | Model | Accuracy | Precision | Recall | F1_Score |
| --- | --- | --- | --- | --- | --- |
| IAV Subtype 5 | LR | 0.9677 | 0.9672 | 0.9677 | 0.9672 |
|  | SVM | 0.9565 | 0.9517 | 0.9565 | 0.9462 |
|  | RF | 0.9774 | 0.9776 | 0.9774 | 0.9757 |
|  | DNABERT2 | 0.9578 | 0.9356 | 0.9578 | 0.9461 |
|  | MegaDNA | 0.9785 | 0.9788 | 0.9785 | 0.9785 |
|  | Influ_BERT | **0.9899** | **0.9901** | **0.9899** | **0.9900** |
| IAV Subtype 10 | LR | 0.9490 | 0.9473 | 0.9490 | 0.9474 |
|  | SVM | 0.9210 | 0.9038 | 0.9210 | 0.9006 |
|  | RF | 0.9533 | 0.9546 | 0.9533 | 0.9444 |
|  | DNABERT2 | 0.9114 | 0.9267 | 0.9114 | 0.9090 |
|  | MegaDNA | 0.9613 | 0.9599 | 0.9613 | 0.9601 |
|  | Influ_BERT | **0.9756** | **0.9753** | **0.9756** | **0.9754** |
| H/N Subtype 10 | LR | 0.9880 | 0.9918 | 0.9880 | 0.9894 |
|  | SVM | 0.9820 | 0.9724 | 0.9820 | 0.9764 |
|  | RF | 0.9880 | 0.9844 | 0.9880 | 0.9844 |
|  | DNABERT2 | 0.9815 | 0.9735 | 0.9815 | 0.9762 |
|  | MegaDNA | 0.9796 | 0.9770 | 0.9796 | 0.9796 |
|  | Influ_BERT | **0.9963** | **0.9960** | **0.9963** | **0.9961** |

Note that “IAV Subtype 5” and “IAV Subtype 10” refer to recognition tasks involving five and ten IAV subtypes, respectively, without distinguishing specific genomic segments; whereas “H/N Subtype 10” denotes the recognition of ten IAV subtypes based solely on the HA and NA segments.

In terms of F1-score **Figure 3a-b**, Influ-BERT demonstrated a significant advantage compared with the other five models. In the five-class(five-subtype) identification task, the F1-scores of influ-BERT achieved 97% and 94% for H5N1 and H9N2, respectively, which is significantly improved when compared to the other models. In the ten-class(ten-subtype) task, Influ-BERT also outperformed almost all other models on all low-frequency subtypes, especially for H5N1, H9N2, H5N8, and H7N9. Notably, in the H5N8 subtype identification task, most other models failed to recognize this subtype, while Influ-BERT achieved an F1-score of 87%, providing strong evidence for the effectiveness of Influ-BERT. Furthermore, Influ-BERT demonstrated consistent performance between recall and F1-score metrics, while exhibiting superior capability in low-frequency subtype identification compared to other benchmark models.

**Figure 3.**
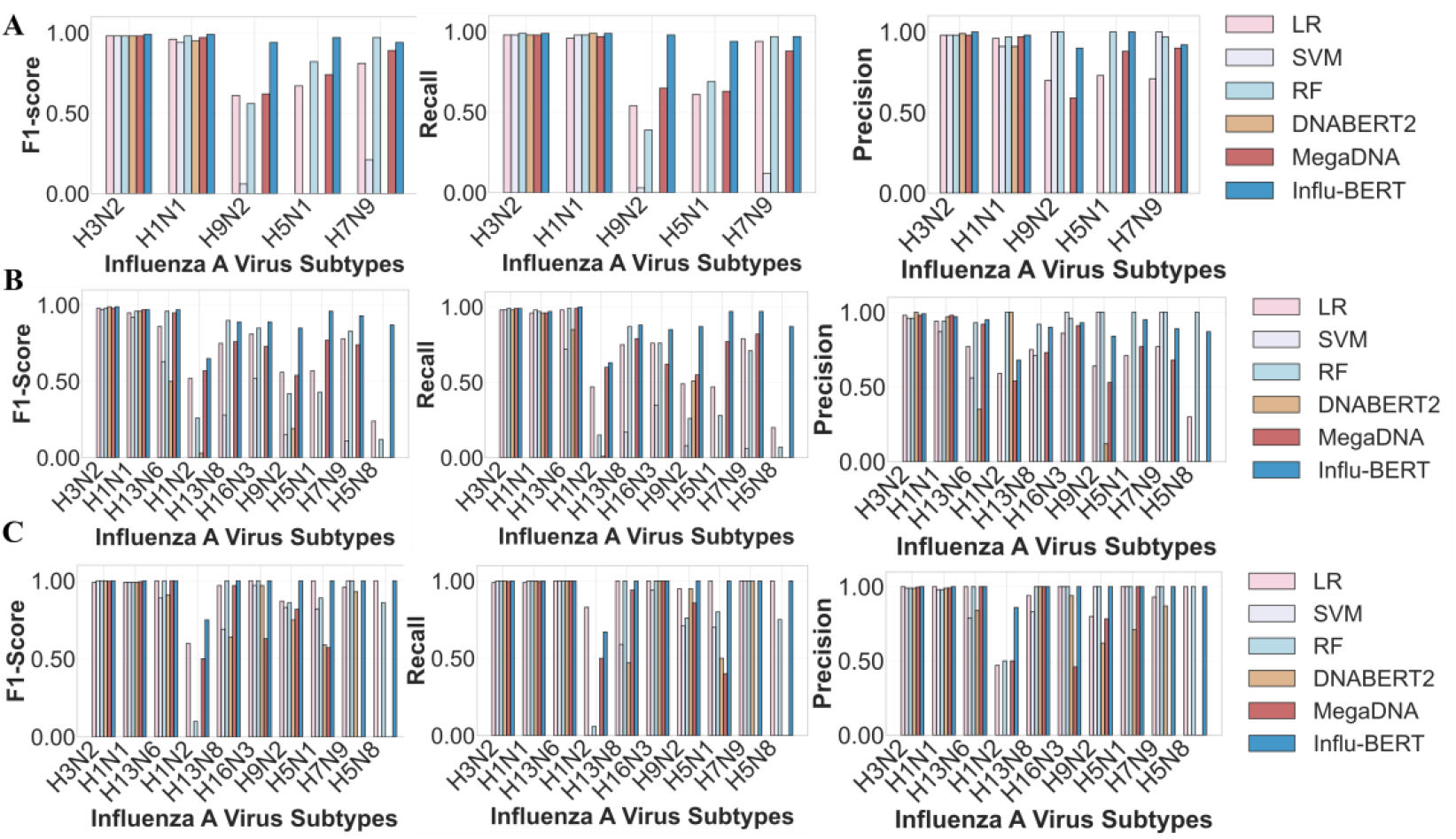
Experimental results of subtyping identification tasks. a) Performance evaluation across five common subtypes without distinguishing specific genomic segments; b) Performance evaluation across ten subtypes without distinguishing specific genomic segments; c) Performance evaluation across ten subtypes using only the HA and NA segments.

In terms of precision, common subtypes like H1N1 and H3N2 achieved high precision for all models. Influ-BERT did not consistently achieve the best performance across all subtypes. In the five-class identification task, Influ-BERT demonstrated superior performance only on the H3N2 and H1N1 subtypes compared to other models, while in the ten-class identification scenario, its advantage was solely observed for the H13N6 subtype. This phenomenon may stem from the significant under-representation of low-frequency subtypes in the dataset. Such class imbalance compels comparative models to adopt more conservative prediction strategies, whereby samples are classified as low-frequency subtypes only when the prediction confidence exceeds a predetermined threshold.

To further address application scenarios that require a focus on specific genomic segments—such as vaccine design—we developed an alternative subtyping workflow. This workflow was evaluated on a ten-subtype classification task. As shown in **Figure 3c**, Influ-BERT outperformed other competing models in this subtype identification (**Table 1**), demonstrating a clear advantage in identifying H1N2 and H9N2 with respect to F1-score. Interestingly, the traditional machine learning model LR also achieved robust performance when limited to the HA and NA segments alone, underscoring the discriminative power of these key genomic regions. Nevertheless, Influ-BERT maintained an overall relative advantage, exhibiting superior performance not only in F1-score but also in recall and precision metrics (**Table 1**), which highlights its effectiveness in scenarios where subtype identification depends primarily on HA and NA sequences. In addition, we further evaluated the model’s ability to distinguish between different types of influenza viruses (see Supplementary Table S2).

In summary, for the scenario without distinguishing specific genomic segments, these experiments found that directly applying the cross-species pre-trained DNABERT-2 model in its original form yielded poor performance in recognizing low-frequency subtypes, and traditional machine learning models also underperformed on low-frequency subtype identification. It is noteworthy that, when compared to conventional machine learning methods, DNABERT-2 exhibited inferior performance, further highlighting the limitations of existing methods in handling extremely imbalanced class distributions. However, Influ-BERT (the modified DNABERT-2 model), after optimization using the two-stage training strategy proposed in this study, achieved significant improvements in identifying both common and low-frequency subtypes, and exhibited consistent improvements across both five-class and ten-class tasks. This demonstrates that Influ-BERT retains a relative advantage in low-frequency subtype identification even as the task scale expands. In the scenario where only the HA and NA segments were used, the performance of the original DNABERT-2 model remained limited. Given that the HA and NA segments are key determinants of IAV subtype combinations, the overall model performance in this experiment was relatively satisfactory, with Influ-BERT outperforming other methods and achieving the highest identification performance as measured by the F1-score.

### Performance Evaluation for Extended Downstream Tasks

To further validate the performance of Influ-BERT, we extended its application to three additional down-stream tasks: identification of common respiratory viruses, identification of IAV genomic segments and functional genes, and prediction of IAV pathogenicity.

In the first task, leveraging the domain-adapted pretrained model, we identified four common respiratory viruses—SARS-CoV-2, rhinovirus, RSV, and influenza virus. As summarized in Table 2, Influ-BERT achieved superior performance compared to other baseline models. We further visualized the embeddings derived from Influ-BERT (**Figure 4a**), which exhibited clear spatial separation between virus types. Applying the K-means clustering algorithm to these embeddings revealed well-defined cluster structures, confirming the model’s ability to learn discriminative sequence representations.

**Table 2.**
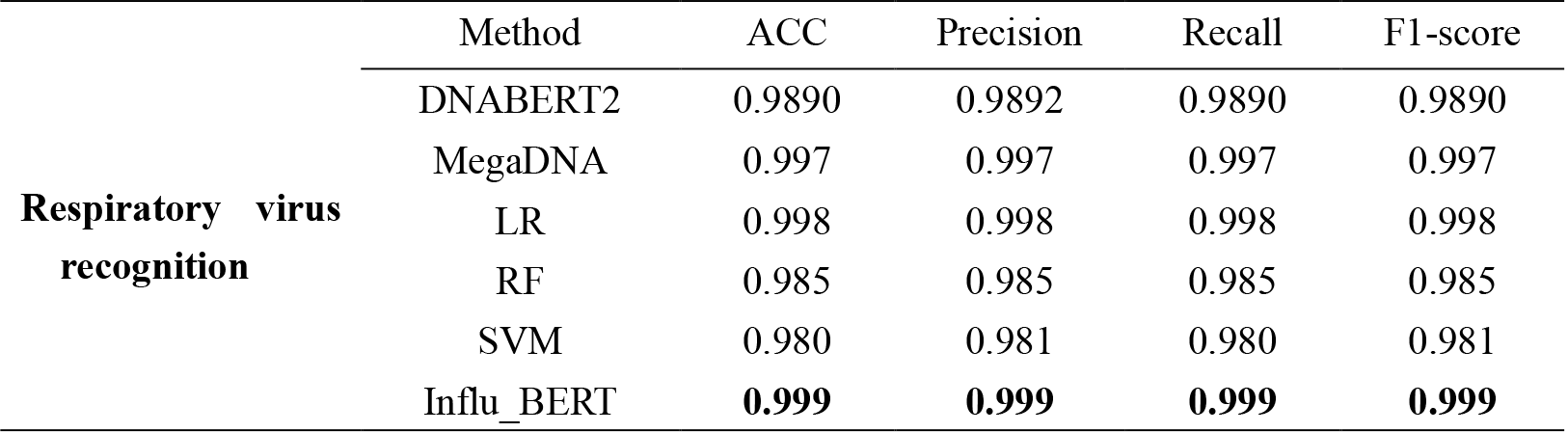
Influ-BERT was compared with baseline models on respiratory virus recognition.

|  | Method | ACC | Precision | Recall | F1-score |
| --- | --- | --- | --- | --- | --- |
| **Respiratory virus recognition** | DNABERT2 | 0.9890 | 0.9892 | 0.9890 | 0.9890 |
|  | MegaDNA | 0.997 | 0.997 | 0.997 | 0.997 |
|  | LR | 0.998 | 0.998 | 0.998 | 0.998 |
|  | RF | 0.985 | 0.985 | 0.985 | 0.985 |
|  | SVM | 0.980 | 0.981 | 0.980 | 0.981 |
|  | Influ_BERT | **0.999** | **0.999** | **0.999** | **0.999** |

**Figure 4.**
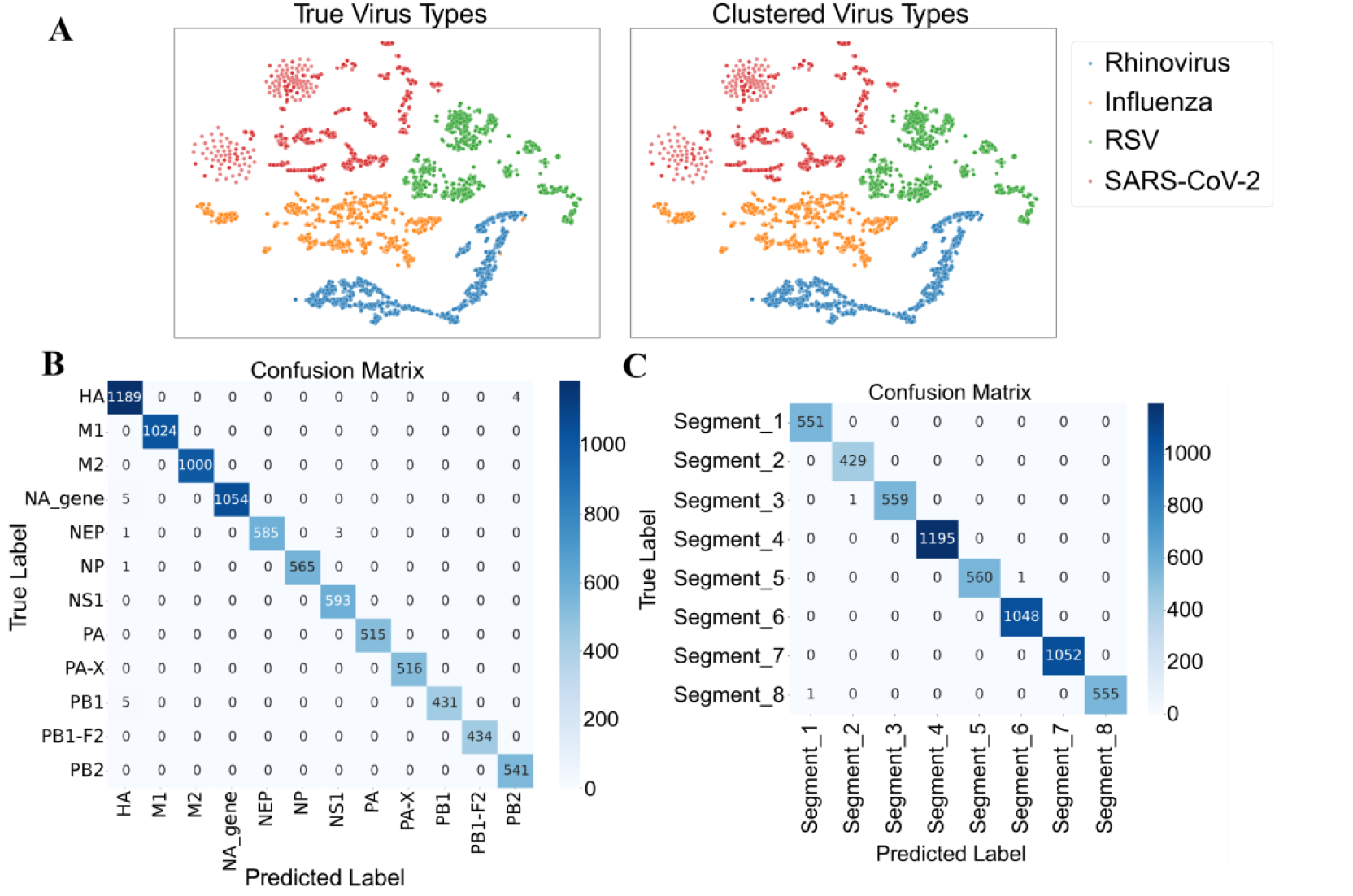
Experimental results of extended tasks. a) Embeddings of diverse respiratory virus sequences, extracted using Influ-BERT, were visualized to examine their distribution patterns; these embeddings were subsequently clustered using the K-means algorithm; b) Confusion matrix for functional gene prediction; c) Confusion matrix for gene segment prediction.

In the second task, we applied Influ-BERT to the identification of influenza virus genomic segments and functional genes. As shown in **Figure 4b**, Influ-BERT accurately identification almost all genomic segments, with only three misclassifications—occurring in Segment 1, Segment 2, and Segment 6— highlighting its robust segment identification capability. The functional gene identification results, presented in **Figure 4c**, show that while the HA gene exhibited relatively more prediction errors compared to other genes, Influ-BERT still achieved excellent overall performance on this task.

In the third task, we addressed the prediction of IAV pathogenicity. Given the limited availabilitity of labeled pathogenicity data in real-world scenarios, this task emphasized evaluating the model’s ability to learn from small datasets. Specifically, we designed two experiments: one with 200-samples and another with 400-samples. As summarized in Table 3, in the 200-samples experiment, Influ-BERT achieved an F1 score of 79.5%, outperforming the second-best model (Random Forest) by 4% and excelling across all evaluation metrics. With increased data in the 400-samples setting, Influ-BERT’s F1 score further improved to 83.3%, maintaining a significant advantage over other models. Notably, DNABERT-2 showed degraded performance in the 400-samples setting compared to the 200-samples setting. These results demonstrate that Influ-BERT possesses strong representation learning capabilities and generalizes effectively when labeled data is scarce.

**Table 3.** Influenza BERT was compared with baseline models on pathogenicity prediction.

|  | Method | ACC | Precision | Recall | F1-score |
| --- | --- | --- | --- | --- | --- |
| 200-samples | DNABERT2 | 0.632 | 0.645 | 0.632 | 0.621 |
|  | MegaDNA | 0.506 | 0.256 | 0.506 | 0.340 |
|  | LR | 0.747 | 0.747 | 0.747 | 0.747 |
|  | RF | 0.756 | 0.765 | 0.756 | 0.755 |
|  | SVM | 0.506 | 0.505 | 0.506 | 0.500 |
|  | Influ_BERT | **0.796** | **0.802** | **0.796** | **0.795** |
| 400-samples | DNABERT2 | 0.564 | 0.567 | 0.564 | 0.562 |
|  | MegaDNA | 0.561 | 0.672 | 0.561 | 0.466 |
|  | LR | 0.781 | 0.781 | 0.781 | 0.781 |
|  | RF | 0.787 | 0.799 | 0.787 | 0.785 |
|  | SVM | 0.572 | 0.574 | 0.572 | 0.571 |
|  | Influ_BERT | **0.833** | **0.836** | **0.833** | **0.833** |

200-samples: the experiment comprised 100 high-pathogenicity and 100 low-pathogenicity samples.

### Interpretability Analysis by Perturbation

To investigate how the model captures informative features within influenza sequences, we designed a masking-based sliding window perturbation analysis. By systematically masking input sequences and observing variations in the model’s predictions, we reveal differences in feature attention patterns between DNABERT-2 and Influ-BERT (**Figure 5**), providing preliminary insights into the latter’s improved performance in low-frequency subtype identification. Using three influenza viral genome sequences of the H5N1 subtype as example (**Figure 5**), both models exhibited varying degrees of attention to the initial nucleotide regions of the sequences. However, Influ-BERT demonstrated relatively consistent overall attention distribution patterns and consistently attended to specific genomic regions near positions 650-700 nt, 1100 nt, 1500 nt, and 1750 nt, indicating robust global modeling capabilities. In contrast, DNABERT-2 displayed irregular and substantially divergent attention distributions across sequences of the same subtype (**Figure 5**), which correlates with its failure to recognize all three sequences. We further conducted perturbation analyses on three subtypes H13N8 and H3N2. The results demonstrate patterns essentially consistent with those observed for H5N1 (**Figure S1-S2**). These findings suggest that Influ-BERT could identify discriminative sequence features that may underlie its superior robustness and accuracy in subtype identification. These identified features or key regions may have practical implications for the model’s decision-making process.

**Figure 5.**
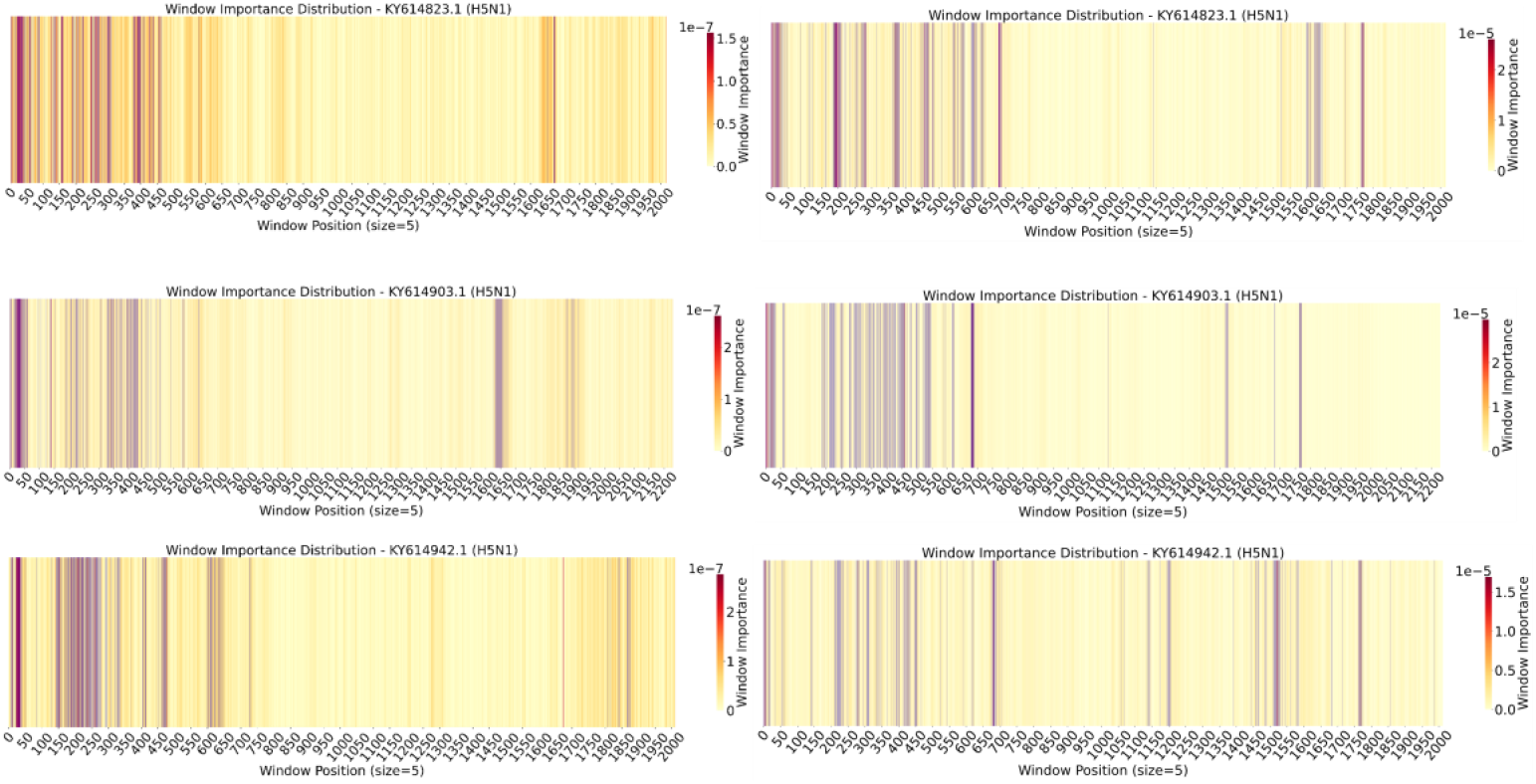
Illustration of perturbation analysis for three randomly selected H5N1 sequences. The left panel shows the results of DNABERT-2, while the right panel corresponds to Influ-BERT.

## Conclusion and Discussion

Although masked pre-trained models such as DNABERT-2, trained on cross-species datasets, and generative models like MegaDNA have demonstrated excellent performance across various biological sequence analysis tasks, they exhibit notable limitations in capturing knowledge representations specific to influenza virus sequences. These constraints stem primarily from the highly long-tailed distribution of influenza virus data, relatively short sequence lengths, and frequent genetic mutations. Notably, in our study, DNABERT-2 underperformed compared with most traditional machine learning models across four downstream tasks.

To address these challenges, we proposed a domain-adaptive pretrained model, Influ-BERT. In the subtype identification task, Influ-BERT demonstrated robust effectiveness even without explicitly distinguishing genomic segments, achieving substantial improvements in both F1-score and recall, particularly for low-frequency subtypes. When trained exclusively on the HA and NA segments, Influ-BERT still outperformed other models. Furthermore, we extended its application to tasks including common respiratory virus classification, IAV gene fragment and functional gene identification, and IAV pathogenicity prediction. Across all these tasks, Influ-BERT consistently achieved superior performance, underscoring its powerful representational capacity for influenza genomic sequences.

From a biological perspective, IAV subtype determination is primarily governed by the HA and NA segments. Nevertheless, AI-based approaches can accurately identify subtypes without explicit segments for two main reasons. First, co-evolutionary relationships among viral gene segments allow discriminative information related to HA and NA to be implicitly captured in other segments [37]. This segment-agnostic identification strategy is particularly advantageous in rapid detection scenarios or when limited sequencing depth leads to missing HA and NA segments. In contrast, for applications that require high accuracy or target specific genomic segments, such as vaccine design, a segment-specific approach is more suitable. This workflow can also be integrated with segment identification models to enable the rapid retrieval of HA and NA segments from large-scale sequence datasets, thereby facilitating subsequent subtype identification processes. Second, the model is capable of learning statistical patterns from sequence data and mapping inputs directly to subtype labels.

Despite the encouraging progress achieved by Influ-BERT in influenza virus sequence recognition, some limitations remain. First, the current model relies on supervised closed-set learning during fine-tuning, which limits its ability to recognize novel subtypes absent from the training data. Future work will incorporate confidence and uncertainty estimation methods to extend the model toward open-set identification. Second, since Influ-BERT was trained solely on DNA sequences, it inherently lacks the ability to capture protein-level structural features. We plan to integrate multi-modal learning approaches to enhance its capacity to interpret protein structural information in future studies.

## Key Points

- We developed an influenza-specific corpus and proposed Influ-BERT using a two-stage pretraining strategy, significantly enhances the model’s capability to identify low-frequency subtypes and overcomes the limitations of general-purpose models when dealing with long-tailed data distributions, short sequence lengths, and high mutation rates.
- We further applied Influ-BERT to three downstream tasks: common respiratory virus identification, IAV gene fragment and functional gene identification, and IAV pathogenicity prediction. Experimental results demonstrate that Influ-BERT achieves consistently superior performance across all tasks, highlighting its powerful representational capacity.
- We introduced a sliding-window perturbation analysis to enhance model interpretability. This method enables the visualization of subtype-discriminative regions and uncovering key attention patterns essential for accurate influenza subtype identification.

## Code Availability

Source code is written in PyTorch and available at https://github.com/oooo111/Influenza-BERT and https://huggingface.co/rongye1/Influenza_BERT under the MIT license.

## Data Availability

All genome sequences were downloaded from https://github.com/oooo111/Influenza-BERT/tree/main.

## Acknowledgements

We thank Yiji Yang for valuable assistance and insightful discussions.

## Funding

This work was supported by the National Key Research & Development Program of China (2025YFF1207901 to S.H.S., 2024YFC2311303 to Z.Z., 2023YFC2604400 to F.C.), the Key Collaborative Research Program of the Alliance of National and International Science Organizations for the Belt and Road Regions (ANSO-CR-KP-2022-09 to S.H.S.), the National Natural Science Foundation of China (32270718 to S.H.S.). ATRV is supported by FAPERJ E-26/201.046/2022 and CNPq (CNPq) 307145/2021-2

## Supplementary Figures Legend

**Supplementary Table S1**. Assigned pathogenicity labels using a label-mapping dictionary.

**Supplementary Table S2**. Baseline comparison for distinguishing the four types of influenza viruses.

**Supplementary Figure S1 H3N2 Sensitivity Analysis**. Influ-BERT demonstrates relatively consistent sensitivity patterns across sequences, whereas DNABERT-2 exhibits greater variability. Notably, Influ-BERT consistently highlights region-specific sensitivity at positions 1300–1350 nt, 1200 nt, 1050–1100 nt, and 350 nt, a pattern not observed in DNABERT-2.

**Supplementary Figure S2 H13N8 Sensitivity Analysis**. Influ-BERT demonstrates a higher consistency in sensitivity distribution, whereas DNABERT-2 shows almost no discernible pattern. DNABERT-2 exhibits a more widespread sensitivity across regions, making it difficult for the model to identify important areas. In contrast, Influ-BERT reveals a more systematic distribution of sensitive regions, indicating that sequences with such a distribution are more likely to be classified as the H13N8 subtype.

## Notes

### Competing Interest Statement

The authors have declared no competing interest.

### Summary of Updates

This version of the manuscript has been revised to include additional downstream task evaluations and to improve the writing.

## References

1. Nair, Harish, et al. “Global burden of respiratory infections due to seasonal influenza in young children: a systematic review and meta-analysis.” The Lancet 378.9807 (2011): 1917–1930.

2. Iuliano, A. Danielle, et al. “Estimates of global seasonal influenza-associated respiratory mortality: a modelling study.” The Lancet 391.10127 (2018): 1285–1300.

3. Li, Xiuli, et al. “Packaging signal of influenza A virus.” Virology Journal 18.1 (2021): 36.

4. Taubenberger, Jeffery K., and John C. Kash. “Influenza virus evolution, host adaptation, and pandemic formation.” Cell host & microbe 7.6 (2010): 440–451.

5. Peacock, Thomas P., et al. “The global H5N1 influenza panzootic in mammals.” Nature 637.8045 (2025): 304–313.

6. Zhang, Jiahao, et al. “H9N2 avian influenza viruses: challenges and the way forward.” The Lancet Microbe 4.2 (2023): e70–e71.

7. Shi, Jianzhong, et al. “Rapid evolution of H7N9 highly pathogenic viruses that emerged in China in 2017.” Cell host & microbe 24.4 (2018):

8. Eisfeld, Amie J., Gabriele Neumann, and Yoshihiro Kawaoka. “Influenza A virus isolation, culture and identification.” Nature protocols 9.11 (2014): 2663–2681.

9. Lee, Ming-Shiuh, et al. “Identification and subtyping of avian influenza viruses by reverse transcription-PCR.” Journal of virological methods 97.1-2 (2001): 13–22.

10. Trombetta, Claudia Maria, et al. “Overview of serological techniques for influenza vaccine evaluation: past, present and future.” Vaccines 2.4 (2014): 707–734.

11. Gauger, Phillip C., and Amy L. Vincent. “Serum virus neutralization assay for detection and quantitation of serum neutralizing antibodies to influenza A virus in swine.” Animal Influenza Virus: Methods and Protocols. New York, NY: Springer US, 2020. 321–333.

12. Ghedin, Elodie, et al. “Large-scale sequencing of human influenza reveals the dynamic nature of viral genome evolution.” Nature 437.7062 (2005): 1162–1166.

13. Patrono, Livia V., et al. “Archival influenza virus genomes from Europe reveal genomic variability during the 1918 pandemic.” Nature communications 13.1 (2022): 2314.

14. Poon, Art F. Y. “Prospects for a sequence-based taxonomy of influenza A virus subtypes.” Virus Evolution 10.1 (2024): veae064.

15. Aksamentov, Ivan, et al. “Nextclade: clade assignment, mutation calling and quality control for viral genomes.” Journal of open source software 6.67 (2021): 3773.

16. Humayun, Fahad, et al. “Computational method for classification of avian influenza A virus using DNA sequence information and physicochemical properties.” Frontiers in Genetics 12 (2021): 599321.

17. Cacciabue, Marco, and Débora N. Marcone. “INFINITy: A fast machine learning-based application for human influenza A and B virus subtyping.” Influenza and Other Respiratory Viruses 17.1 (2023): e13096.

18. Borkenhagen, Laura K., Martin W. Allen, and Jonathan A. Runstadler. “Influenza virus genotype to phenotype predictions through machine learning: a systematic review: computational prediction of influenza phenotype.” Emerging microbes & infections 10.1 (2021): 1896–1907.

19. Vaswani, Ashish, et al. “Attention is all you need.” Advances in neural information processing systems 30 (2017).

20. Theodoris, Christina V., et al. “Transfer learning enables predictions in network biology.” Nature 618.7965 (2023): 616–624.

21. Sapoval, Nicolae, et al. “Current progress and open challenges for applying deep learning across the biosciences.” Nature Communications 13.1 (2022): 1728.

22. Peng, Cheng, et al. “ViraLM: empowering virus discovery through the genome foundation model.” Bioinformatics 40.12 (2024): btae704.

23. Consens, Micaela E., et al. “Transformers and genome language models.” Nature Machine Intelligence (2025): 1–17..

24. Nguyen, Eric, et al. “Sequence modeling and design from molecular to genome scale with Evo.” Science 386.6723 (2024): eado9336.

25. Ji, Yanrong, et al. “DNABERT: pre-trained Bidirectional Encoder Representations from Transformers model for DNA-language in genome.” Bioinformatics 37.15 (2021): 2112–2120.

26. Zhou, Zhihan, et al. “Dnabert-2: Efficient foundation model and benchmark for multi-species genome.” arXiv preprint 2306.15006 (2023).

27. Shao, Bin, and Jiawei Yan. “A long-context language model for deciphering and generating bacteriophage genomes.” Nature Communications 15.1 (2024): 9392.

28. Hou, Xin, et al. “Using artificial intelligence to document the hidden RNA virosphere.” Cell 187.24 (2024): 6929–6942.

29. Krammer, Florian, and Stacey Schultz-Cherry. “We need to keep an eye on avian influenza.” Nature Reviews Immunology 23.5 (2023): 267– 268.

30. Petrova, Velislava N., and Colin A. Russell. “The evolution of seasonal influenza viruses.” Nature Reviews Microbiology 16.1 (2018): 47–60.

31. Caserta, Leonardo C., et al. “Spillover of highly pathogenic avian influenza H5N1 virus to dairy cattle.” Nature 634.8034 (2024): 669–676.

32. Federhen, Scott. “The NCBI taxonomy database.” Nucleic acids research 40.D1 (2012): D136–D143.

33. Breiman, Leo. “Random forests.” Machine learning 45 (2001): 5–32.

34. Pisner, Derek A., and David M. Schnyer. “Support vector machine.” Machine learning. Academic Press, 2020. 101–121.

35. Hosmer Jr, David W., Stanley Lemeshow, and Rodney X. Sturdivant. Applied logistic regression. John Wiley & Sons, 2013.

36. Niu, Shuteng, et al. “A decade survey of transfer learning (2010–2020).” IEEE Transactions on Artificial Intelligence 1.2 (2021): 151–166.

37. Obenauer J C, Denson J, Mehta P K, et al. Large-scale sequence analysis of avian influenza isolates[J]. science, 2006, 311(5767): 1576–1580.

