## Supplementary information for "Influ-BERT: A Domain-Adaptive Genomic Language Model for Advancing Influenza A Virus Research"

### Supplementary materials

| High pathogenicity label | Low pathogenicity label |
|---|---|
| HPAI | LPAI |
| High path | Low path |
| Highly pathogenic | Lowly pathogenic |
| Severe disease | Mild disease |
| High mortality | Low mortality |
| Lethal | Non-lethal |
| Fatal infection | Mild infection |
| Severe infection | Asymptomatic |
| High virulence | Low virulence |
| Severe clinical | Mild clinical |
| Deadly | Mild |
| Death | Moderate symptoms |
| Fatal | Low severity |
| Severe symptoms | |
| High severity | |

**Supplementary Table S1**

Analysis：We retrieved influenza A virus (IAV) sequences from the NCBI database and assigned pathogenicity labels using a label-mapping dictionary.

| Task | Model | Accuracy | Precision | Recall | F1_Score |
|---|---|---|---|---|---|
| FLU typing | DNABERT2 | 0.9874 | 0.9877 | 0.9874 | 0.9868 |
| | MegaDNA | 0.9978 | 0.9978 | 0.9978 | 0.9978 |
| | Influenza BERT | **0.9982** | **0.9982** | **0.9982** | **0.9982** |

**Supplementary Table S2**

Analysis：We conducted a baseline comparison for distinguishing the four types of influenza viruses, based on 20,000 IAV sequences, 20,000 IBV sequences, and all available ICV and IDV sequences.

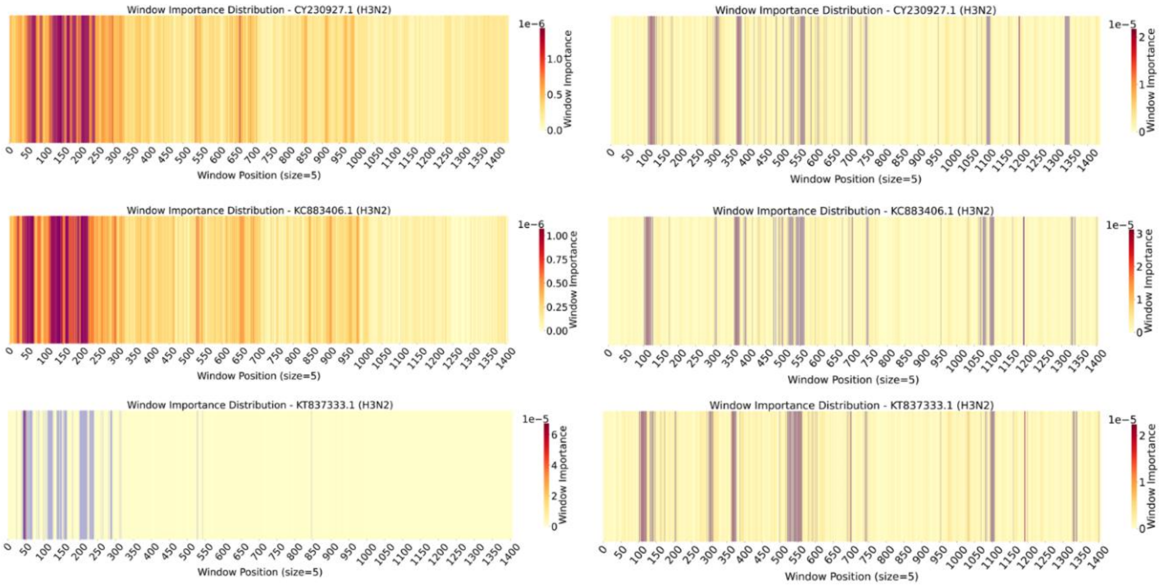

#### Supplementary Figure S1

Analysic: As shown in Figure S1, Influenza BERT demonstrates relatively consistent sensitivity patterns across sequences, whereas DNABERT-2 exhibits greater variability. Notably, Influenza BERT consistently highlights region-specific sensitivity at positions 1300–1350 nt, 1200 nt, 1050–1100 nt, and 350 nt, a pattern not observed in DNABERT-2.

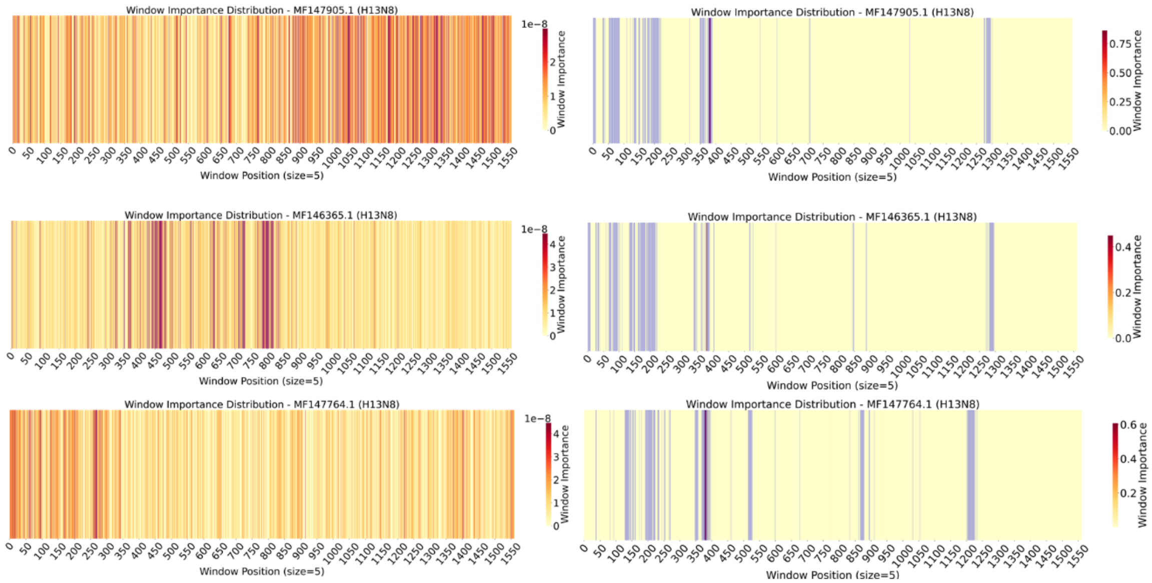


#### Supplementary Figure S2

Analysic: As shown in Figure S2, Influenza BERT demonstrates a higher consistency in sensitivity distribution, whereas DNABERT-2 shows almost no discernible pattern. DNABERT-2 exhibits a more widespread sensitivity across regions, making it difficult for the model to identify important areas. In contrast, Influenza BERT reveals a more systematic distribution of sensitive regions, indicating that sequences with such a distribution are more likely to be classified as the H13N8 subtype.
